## Supplementary Figures for "A single amino acid variant in the variable region I of AAV capsid confers liver detargeting"

**a**

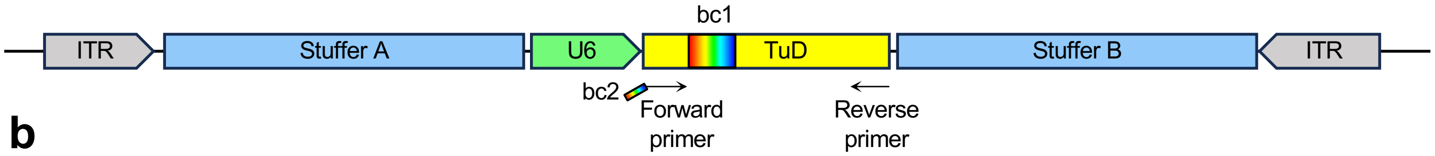


**b**

ctgcgcgctcgctcgctcactgaggccgcccgggcaaagcccgggcgtcgggcgacctttggtcgcccggcctcagtgagcgagcgagcgcgcagagagggagtggccaactccatcactaggggttccttgtagttaatgat
taacccgccatgctacttatctaccagggtaatggggaagtacaataaaaacaaggctttcaagctgataaaggcttctttcccactcatagaggaaattctttaaagtatccccaaaattgctttcctgaggtgttcctcttggtt
aaccgtctagtgattggggacctttgaaaacactcaggattgaagataatggtcttttctcactattggtgatgtaatcagtgttctgaaaataaaatactattttagctagagcgttatagaagtcagtagaatattcagaatttc
atttgcatttctgatcatcaaggtgaccaccactgccagcatgtgcagggtacctgcattgtgtctcttggccacagcgggtccttctgcaggtggagctgtggtctttgcagttgggcgcagctagcatggcaggtacatgagca
tttggaagagcagtcaggacccatgtgctcagcacatgtgttttcttgggcagttggcagaaaccataagtcttttcccaacttttatgtttgggaagagagagaaagaagccaaatttgctatacgtttgatgttattaaggttgt
catttgaataagtttttaaaaaatgtataacataataactggattgatacaagcatgctgaaagataaactaataactgaacatatccgttaatttttttggtatttgtagtgaaaacataattgtaattgaaaacaacattactg
aagagggaaaaatgatttttatgcctcaagcaaaacacacacatatacacaaaaaattaaagacttaagcttttttttttctttccaagaacatctggtttacagtatgctaaaagcaaatttggattaacctagaaattaagtg
tgtgtcatatacaagccattgattaaaccatattaaaatatttagacctcatggaaaggtataaaactctccaccatgtacagatttttatttttttgtaaggaaagtaaacaaaactatttcaatgaatcctgtgtcaaaaaaatt
aaaataaatctgtgtgatgtcatctgcgtctgccaacctgcccttccaaagagacgcaagatggaagctttcagtgcttgtcctgaccacccagcaacaggtctttaagcctgaggacagtcattccttacacttttattctccaat
ttctttgctttgtgtaggactttacgggcatctatgttcattcttaaaatatttcgtactaatataaaaatcgtggacctgagtccaagccacaaaccctgaagcaagaatttggtcctgatcttgagtcttctttcacctgcagagcc
ctccccggactctcaggaccataggctgtccaggcactcctgattctgaagccacgggaagtcccaatttcagccattgttggtgagaaacaatccctctttaagagctgcccataagggcaacctgtcccaagaggcttccgat
agcatctgcgtcctcttctgcatgttgtcacccacgaaggaggcctgcgtggcacagagcccacagtgccatatttctgaagcagtggaaggaattgctctgggtgacgccggtggaaggaattgctctccgctgaccctagca
atggatgctgagccaggagcaagtgagaacaaggtcccacgagatcagccggagggcatggggaataagcttctgtcccttcaaggaggccaagtaagataagtttctagttctgtcacattcacagacatctagttcatgac
acagagccttgctttttccaccaggaataaaataccccgatcagcagcttgtgataagttctccccacctgctccctccagcatcccaggagagaacgttaactacacaccttgtttaatgcaaagtcttataatatccatgtgtct
tagaaaataaaattgtagagaccaagaagtgcaaataatcaaattatttaataggaaatttgtgttaggaattccccagtggaaagacgcgcaggcaaaacgcaccacgtgacggagcgtgaccgcgcgccgagcgcgcg
ccaaggtcgggcaggaagagggcctatttcccatgattccttcatatttgcatatacgatacaaggctgttagagagataattagaattaatttgactgtaaacacaaagatattagtacaaaatacgtgacgtagaaagtaat
aatttcttgggtagtttgcagttttaaaattatgttttaaaatggactatcatatgcttaccgtaacttgaaagtatttcgatttcttgggtttatatatcttgtggaaaggacgcgggatccgacggcgctaggatcatcaac**NNN**
**NNNNN**acgaatctctatatcatcaagtattctggtcacagaatacaacaagccacaacgaatctctatatcatcaagatgatcctagcgccgtcttttttaagctgtcccggatccgacccaggacctaactacgaaagcag
ttttacgtctaaaacacagagatgttttgttactcaagctatacggcatgttgcctttccctgacaaggcagtgtgcctgagaatggatgggatataattattttttttaattccagaaaatttgcttgatgttcataaataatcgatt
caatcactttagcaaaagggaaatgttaaaatgatggaaaattctgttcacctagacttgtctgcagtctttaaatcattctccactccatccctcttgctgacattcggagatgatttcaattaagggcctttgacaaagtactat
agtcatttaaaagatggctttaaagagttcagaaatcccaatagttctccatatcaagagggtgcactataaaattgtaaatgagtgcatgcactggctggggttataaagtgccgttgtttttctggtttgacatttcatggattc
caagtgggcctagcttagtggtgcaacataaatcacggaacggatctggatattgtaaaacttgaatatagaagcattttagaggctgtgaagttaacagtaatcagtaagtattctaatagtatgagttactgctaacattata
attatagagtaaaataataatttttataggcaaggagcactgaaaactaatggtcaggatgattataatagctatgaattaatagatcagttgaaattgtgatgactattttaataatgagactgaaatatcgtttgtgttaaaac
agtgagtctaagagcataatcaaccctcttttgtcaggaagaaacaattccttttaactccaaggtcaccagaatgatcctagagttgtattgtggttactctttctccttggaaatgattcttccgttctcctttgctctgtagagtc
ctattcctctacctcatcaggtaggtctgtgtgtgtgagaatgtttgttctcctttgccctgtagagtcctattcgtctaccttgtcgtgtaggtttgtgtgtgggcgagtatttgttctcctttgccctgtagagtcctattcctctacctc
gtcgtgtaggtttccgtgtgggggaatgttgctcccatatcagaacagaagcaccttgcagggggaaggcttgtttccttaaggtctccgtatcctaaattaccacaaaggtgctgaaaatcagcaatcaaggcttgagttgcat
gaactgagctagttacactgcattacagtgactctgcagacaccactggacacagcaaagggccagaagacagcaagccttggccatagccgctgtcatcattcactccccacctcctaggagaaggaagatggaaacaag
tcaatgagttcataaaacagaaacccaggagtcattgcatcactccttatggctcagtggctcagattcactttataaaaacatactaatttcaatttcacatggtatgattcaaagagtcagaaattcatcgtctatcaagtggt
ctgaatgttatcctgcagaattttctctgaagatgaactggttcatccctagggctgtgcaatagtgtctcaatatccaatcactccccactatgtgaattaggccatatgttcttcaccggaacatttgaagaactcttgttgagat
tgtgcagttttagacactgggtcatcaagataaaagacacagaagctctgagaacacacaacttattttcacttgtccaaaataatcatcggctagtaagcttttattcattcatttttaaattgtcattgaaaaatgtatcaactt
agttttagaggaaaatcagtcatgtttgttaaattgatcatctacttgaaaactgtctttctggaaagaagaaaagatgttgctctctgtgcctgcatcctttggtgaggcctggagatgacctgggaaaaacacttggtgaccatt
accctggtagataagtagcatggcgggttaatcattaactacaaggaacccctagtgatggagttggccactccctctctgcgcgctcgctcgctcactgaggccgggcgaccaaaggtcgcccgacgcccgggctttgcccgg
gcggcctcagtgagcgagcgagcgcgcag

**c**

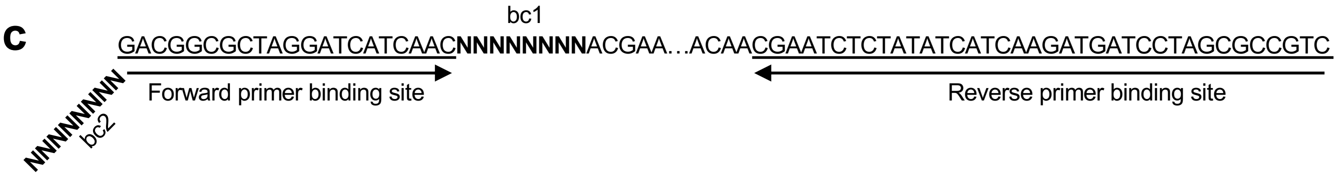


**Supplementary Figure 1. Barcoded construct and PCR design. (a)** Schematics showing the barcoded AAV construct, not drawn to scale. The first barcode (bc1) in the TuD and the second barcode (bc2) in the forward PCR primer are rainbow-colored. Note that each unique bc1 was used to pair with a unique capsid variant during rAAV production, and that bc2 was used in PCR to differentiate among tissue samples. **(b)** The DNA sequence of the construct shown in a. Key elements are color-coded as the scheme shown in a. Gray: ITR. Blue: stuffer. Green: U6 promoter. Yellow: TuD. NNNNNNNN: bc1. **(c)** Binding sites of the forward and reverse PCR primers shown in a. Binding sites are underlined. Note that bc2 is included at the 5' end of the forward primer.

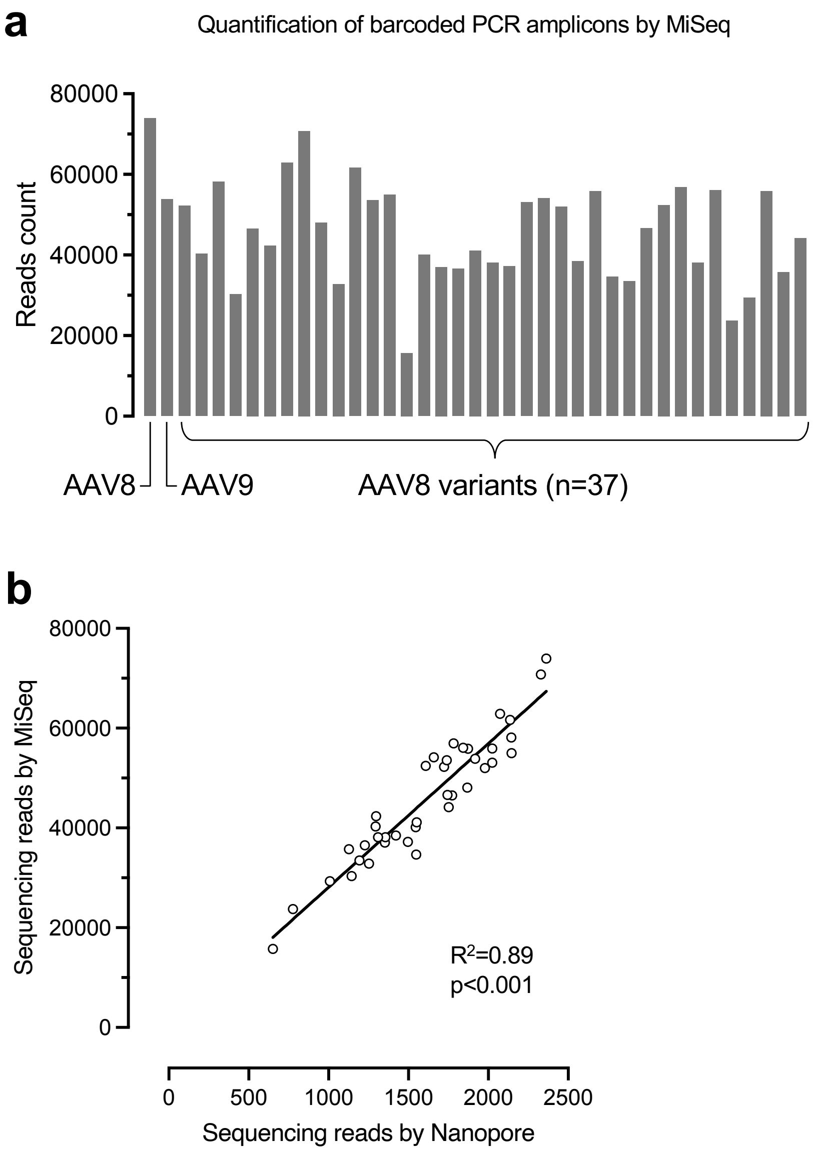


**Supplementary Figure 2. Validation of the relative distribution of barcoded PCR amplicons derived from vector library DNA. (a)** Bar graph showing the count of Illumia sequencing (MiSeq) reads mapped to the unique vector transgene barcodes packaged in AAV8, AAV9, or AAV8 variants. Data were based on one biological repeat. **(b)** Scatter dot plot showing the relationship between the sequencing read counts by nanopore method (x-axis) and MiSeq method (y-axis). The linear regression statistics are shown.

| | Ferret ID | Sex | At rAAV injection | At euthanasia |
|---|---|---|---|---|
| anti-AAV8 | 687721 | M | <1:5 | 1:320 |
| | 687741 | M | <1:5 | 1:320 |
| | 678861 | F | <1:5 | 1:640 |
| anti-AAV9 | 687721 | M | <1:5 | 1:640 |
| | 687741 | M | <1:5 | 1:640 |
| | 678861 | F | <1:5 | 1:640 |

**Supplementary Figure 3. Tabulation of neutralizing antibody titers in ferret serum samples.** Serum samples were collected immediately prior to rAAV injection (at rAAV injection) or immediately prior to euthanasia (at euthanasia). M: male. F: female.

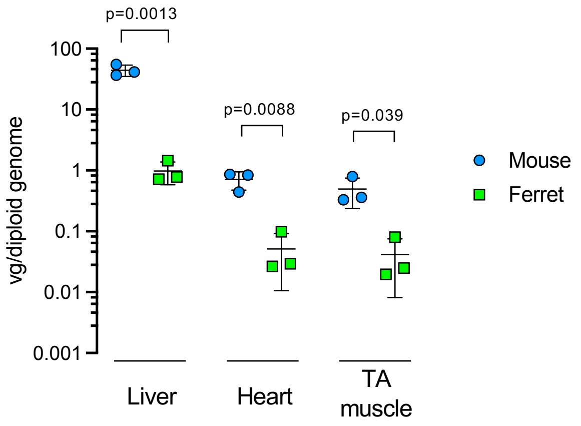


**Supplementary Figure 4. Vector genome abundance in mouse and ferret tissues.** Scatter dot plot showing the vector genome copy number per diploid host genome (vg/diploid genome) in the liver, heart, and tibialis anterior (TA) muscle collected from the mice (blue dots) and ferrets (green dots) treated with the vector library. Each dot represents an individual animal. Data are shown as mean and standard deviation. Statistical analysis is performed using t-test.

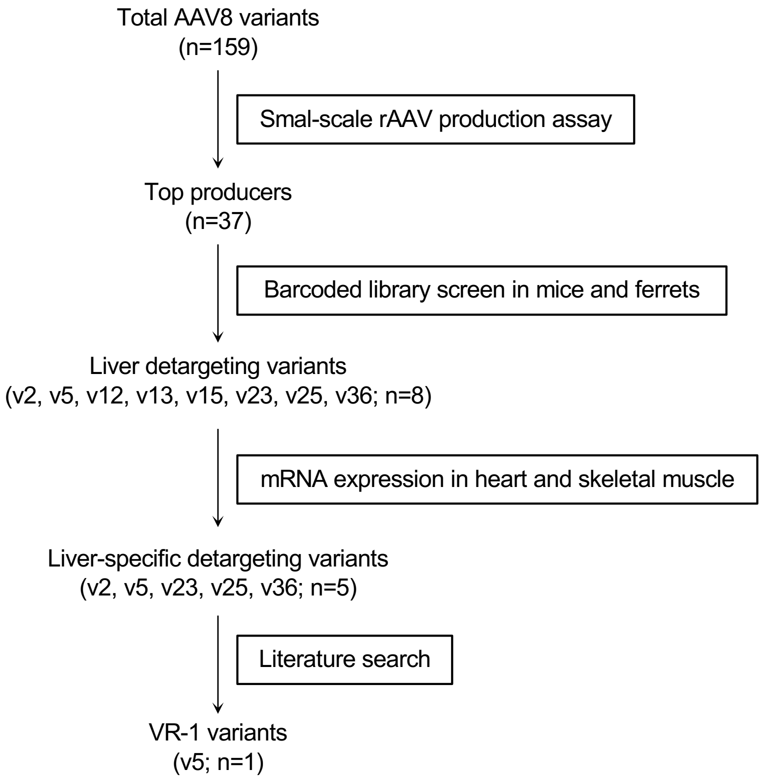


**Supplementary Figure 5. Workflow to select AAV8 variants.**

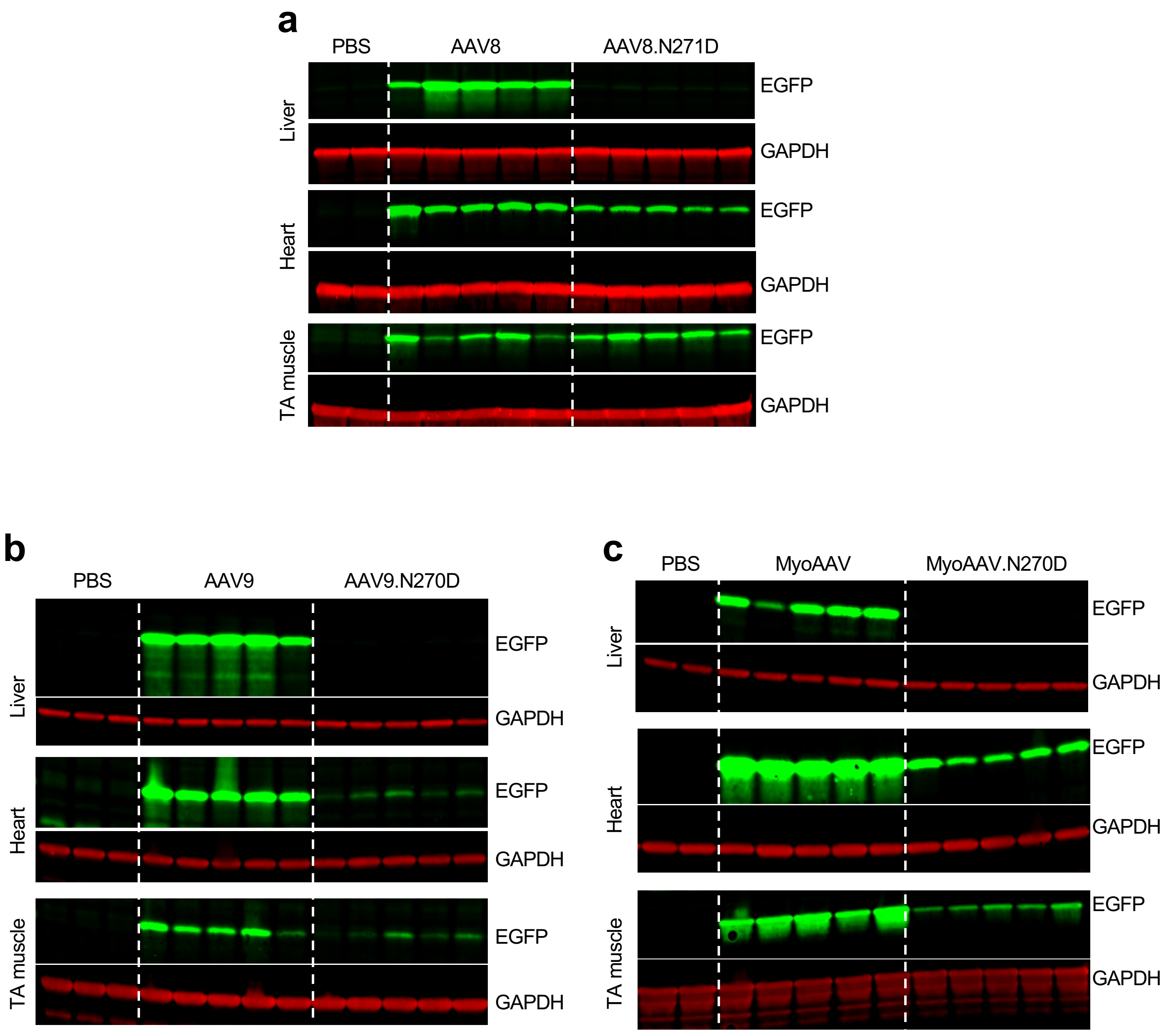


**Supplementary Figure 6. Images of western blotting. (a-c)** EGFP (green) and GAPDH (red) signals in the liver, heart, and tibialis anterior (TA) muscle tissue lysates. Mice were treated with AAV8 or AAV8.N271D vectors (a), AAV9 or AAV9.N270D vectors (b), and MyoAAV or MyoAAV.N270D vectors (c). Mice treated with PBS serve as negative controls. Different treatment groups are separated by dashed white lines to enhance visualization. Each lane represents an individual mouse.

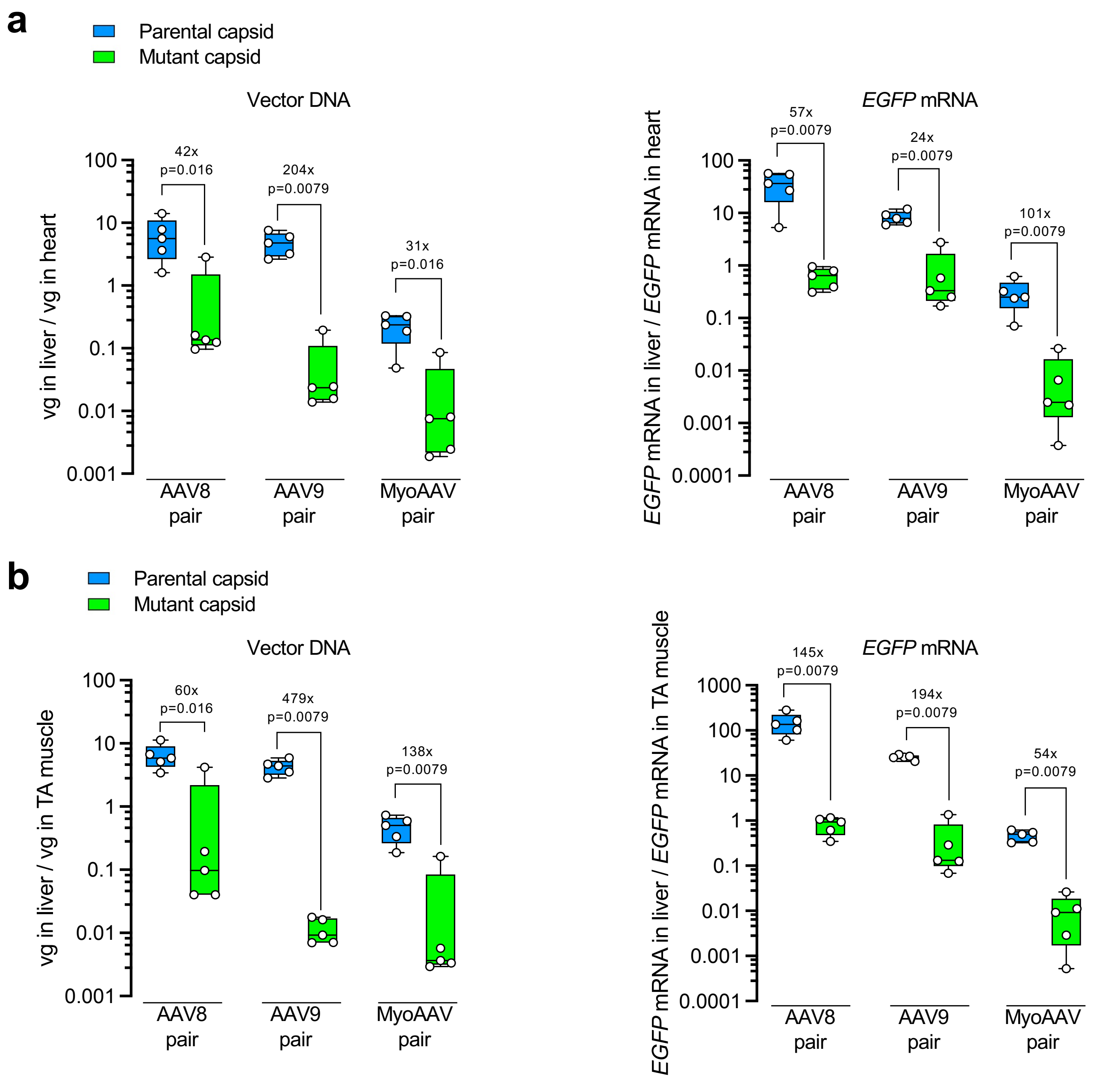


**Supplementary Figure 7. Relative gene delivery and transgene expression.** The levels in liver relative to those in heart (a) and TA muscle (b) in individual mice are calculated using the data shown in Figure 4 and plotted. Each dot represents an individual mouse. The box extends from the first to the third quartiles with the line inside denoting median. The whiskers end at minimum and maximum values. The fold changes of medians and p values are labeled. Statistical analysis is performed using non-parametric Mann-Whitney test.
